# Effects of working memory training on cognitive flexibility, dendritic spine density and long-term potentiation in female mice

**DOI:** 10.1101/2024.07.24.603721

**Authors:** Vasiliki Stavroulaki, Lida-Evmorfia Vagiaki, Orestis Nikolidakis, Maria Zafiri, Maria E. Plataki, Kyriaki Sidiropoulou

## Abstract

Working memory (WM) is a cognitive function that refers to the ability of short-term storage and manipulation of information necessary for the accomplishment of a task. Two brain regions involved in WM are the prefrontal cortex (PFC) and the hippocampus (HPC). Several studies have suggested that training in WM (WMT) can improve performance in other cognitive tasks. However, our understanding of the neurobiological changes induced by WMT is very limited. Previous work from our lab has shown that WMT enhances synaptic and structural plasticity in the PFC and HPC in male mice. In this study, we investigate the effect of WMT on cognitive flexibility and synaptic properties in PFC and HPC in adult female mice. To this end, female adult mice were split into 3 groups: a) naïve which remained in their home cage, b) non-adaptive which learned to alternate the arms in the T-maze but without any delays and c) adaptive which were trained in the delayed alternation task for 9 days. The delayed alternation task was used for WMT. In one cohort, following the delayed alternation task, all mice were tested in the attention set-shifting (AST) task in order to measure cognitive flexibility, and then, the brains were harvested for Golgi-Cox staining to study dendritic spine density. Our results showed that in female mice, there were no differences in performance in the AST among the three groups tested, however, the latency to make a choice was reduced. With regards to dendritic spine density, no significant differences were identified in PFC while increased dendritic spine density was found in the hippocampus of the adaptive group, compared to the naïve group. In a second cohort, acute brain slices were prepared following the delayed alternation task to investigate the synaptic properties in the PFC and the HPC. Evoked field excitatory post-synaptic potential (fEPSP) recordings were performed in either PFC or HPC brain slices. Our results show that tetanic-induced long-term potentiation (LTP) in the PFC was not different among the three training groups. In the HPC, theta-burst induced LTP was significantly increased in the adaptive group also compared to the other two groups. These results reveal both similarities and differences of WMT on cognitive flexibility, dendritic spine density and LTP in females, compared to males.

## Introduction

Working memory (WM) is a fundamental cognitive function that allows the temporary storage and manipulation of information which guides subsequent action (Goldman-Rakic, 1995). The prefrontal cortex (PFC) and the hippocampus (HPC) are two major structures involved in spatial working memory (Pastalkova et al., 2008; Kesner and Churchwell, 2011; Spellman et al., 2015). Both temporally stable and dynamic neuronal activity in the PFC underlies proper WM function (Murray et al., 2017; Wasmuht et al., 2018) in well-trained monkeys and rats (Baeg et al., 2003). Neuronal activity in the PFC is also necessary for successful performance of WM tasks in rodents (Baeg et al., 2003; Kamigaki and Dan, 2017). On the other hand, the HPC also mediates performance in working memory tasks (Avigan et al., 2020; Martins et al., 2019; Wang & Cai, 2006) possibly by representing temporal relationships (Taxidis et al., 2020). Furthermore, the functional connections and communication between PFC and HPC seem to support the encoding phase in spatial working memory tasks (Spellman et al., 2015; Myroshnychenko et al., 2017; Tamura et al., 2017).

Since WM is taxed in a series of other cognitive functions, the effect of working memory training (WMT) on other cognitive tasks has been investigated, yielding both negative and positive effects (Stavroulaki et al., 2020). These conflicting results may stem from our limited understanding of the neurobiological changes that occur in response to WMT. Studies of the effect of WMT on behavior and brain structure and function in rodents are very few (Light et al., 2010; Stavroulaki et al., 2021; Barbelivien et al., 2024). In a previous study, the beneficial effect of WMT on cognitive flexibility as well as synaptic plasticity and dendritic spine density was identified in male mice (Light et al., 2010; Stavroulaki et al., 2021). However, the effect of WMT on cognitive flexibility or neurobiological parameters of PFC and HPC have not been investigated in female mice.

Therefore, the present study aims to fill this gap by investigating the effects of WMT, using the delayed alternation task in the T-maze, on cognitive flexibility, dendritic spine density and LTP in both PFC and HPC.

## Materials and Methods

### Animals

All mice were bred and housed in the Dept of Biology, University of Crete facility. Female C57/B6 mice, aged 5-7 months old were housed in groups (2–4 per cage) and provided with standard mouse chow and water ad libitum, under a 12 h light/dark cycle (light on at 8:00 a.m.) with controlled temperature (24±1°C). All procedures in animals were performed in compliance to the ARRIVE guideline. All animal experimentation protocols approved by the Research Ethics Committee of the University of Crete and obey the European Union ethical standards outlined in the Council Directive 2010/63 EU of the European Parliament on the protection of animals used for scientific purposes.

### Behavioral tasks

To train working memory, we used the delayed alternation task in the T-maze, an apparatus that includes a start arm and two goal arms (45×5cm each). Mice were initially handled by the experimenter for 10 days, and then habituated in the T-maze apparatus, for 2 days. Mice were food-restricted so that the animals maintained 85-90% of their initial weight. Mice from each cage were randomly split into three groups: a) naïve group, b) non-adaptive group and c) adaptive group. Mice in the naïve group remained in their homecage, while mice in non-adaptive and adaptive groups continued training in the T-maze, in the alternation task. All the mice were subject to 10-trial sessions, 3 sessions/day. At the first trial of each session, mice were allowed to freely choose between the right or left goal arms. In the following trials, mice had to alternate the goal arms in order to receive reward, initially with no temporal delay between the trials. Once they reached a predefined criterion for the alternation procedure [i.e., 2 consecutive sessions of ≥70% correct choices (performance)], mice were split into the non-adaptive and adaptive groups. Mice in the non-adaptive group continued to perform the same alternation task for 2 sessions per day. Mice in the adaptive group started the delayed alternation procedure, for which delays were introduced, initially at 10 seconds and increasing by 10 seconds when the criterion for each delay was achieved, for 9 days.

Two days following training, all groups of mice were subjected to the attentional set shifting task (AST), a behavioral task that examines cognitive flexibility (Brown and Tait, 2016). For this task, an open field device is used along with two small identical cups, which contain various substrates, the olfactory cues and food reward. For 2 days, mice habituate to the open-field and they learn to dig for food in the bowl filled with sawdust. During the different experimental stages, mice need to pay attention and respond to the relevant cue (digging medium) and ignore an irrelevant cue (odor), by pairing a food reward with the medium. This association is then reinforced in subsequent stages where the type of digging medium and odor changes, before changed entirely in the final stage. This task contains 7 stages, namely: a) simple discrimination (SD) in which mice differentiate between two different beddings, b) compound discrimination (CD), in which mice still differentiate between two different beddings, but two different olfactory cues are introduced that are not relevant, c) compound discrimination reversal (CDR) in which the bedding and olfactory cues are the same, but the rewarded bedding is switched, d) intradimensional shift 1 (ID-1), in which two different bedding media and olfactory cues are introduced, but one of the beddings is rewarded, e) intradimensional shift 2 (ID-2), in which two different bedding media and olfactory cues are introduced, but one of the beddings is rewarded, f) interadimensional shift 2 reversal (ID-2R), in which the bedding and olfactory cues are the same as in ID-1 but the rewarded bedding is switched, and g) extradimensional shift (ED), in which one of the olfactory cues is rewarded. For mice to successfully complete each stage, they had to complete six consecutive correct trials. The total # of trials required to reach criterion and the latency to first reach the bowl in all trials were recorded for each stage.

### Golgi Cox staining

After completion of the AST task, mice were euthanized, their brains were removed and subjected to Golgi Cox staining. In some mice, technical problems did not allow for successful imaging procedure to take place. This staining remains a key method to study neuronal morphology (Zaqout and Kaindl, 2016). Specifically, mice brains were removed, isolated and placed in a glass vial in a Golgi Cox solution (5% Potassium Dichromate in dH2O, 5% Mercuric Chromate in dH2O, 5% Potassium Chromate in dH2O), that it made and stored 5 days in the dark. The solution is renewed day by day for 10 days. The eleventh day, brains transfer to a 30% sucrose solution in dH2O at 4° C, until the day they were slicing with a vibratome. 150□μm coronal slides of the prefrontal cortex and hippocampus were obtained with a vibratome (VT1000S, Leica Microsystems, Wetzlar, Germany) and placed on gelatin-coated slides. After finishing the sectioning of all samples, the slices kept at 4°C in dark for one day in a humidity chamber. The tissue sections were, then, treated with ammonium hydroxide for 15min and fixed with Kodak Fix solution (30 min per solution). Finally, the tissue was rinsed, dehydrated in graded concentrations of alcohol, incubated in xylene for 5min and coversliped with permount.

One month later, the slices were observed and photographed under a light microscope (NikonPlan Apo 60x/140oil WD 0.21 lenses with a Nikon Eclipse E800 microscope). From each animal and each brain area, we analysed 3-5 sections, and in each section 1-3 neurons from both hemispheres. From each neuron, the number of spines on well-identified secondary dendritic branches (one field of view each) was counted along with the length of each dendritic segment measured. The spines were classified according to the spine type: mushroom, thin (referred as mature) and stubby spines. Images were also obtained from all dendritic segments analyzed and processed using Adobe Photoshop.

### Electrophysiological recordings

In another cohort of mice, 1-5 days following the end of the delayed alternation task protocol, mice were prepared for electrophysiological experiments using the *in vitro* slice preparation. This range (up to 5 days) was similar to the number of days required for mice to perform the AST task in the other group. We did not observe any differences dependent on the day of the recording. The person performing the electrophysiological experiments was blind to the type of behavioral training on the animals. Mice were decapitated under halothane anesthesia. The brain was removed immediately and placed in ice cold, oxygenated (95% O_2_/5% CO_2_) artificial cerebrospinal fluid (aCSF) containing (in mM): 125 NaCl, 3.5 KCl, 26 NaHCO_3_, 1 MgCl_2_ and 10 glucose (pH=7.4, 315 mOsm/l). The brain was blocked and glued onto the stage of a vibratome (Leica, VT1000S, Leica Biosystems GmbH, Wetzlar, Germany). Brain slices (400μm thick) containing either the PFC or the HPC were taken and were transferred to a submerged chamber, which was continuously super-fused with oxygenated (95% O2/5% CO2) aCSF containing (mM): 125 NaCl, 3.5 KCl, 26 NaHCO_3_, 2 CaCl_2_, 1 MgCl_2_ and 10 glucose (pH=7.4, 315mOsm/l) in room temperature. The slices were allowed to equilibrate for at least an hour in this chamber before experiments began. Slices were then transferred to a submerged recording chamber (Scientifica, Inc), which continuously super-fused oxygenated (95% O2/5% CO2) aCSF containing (in mM): 125 NaCl, 3.5 KCl, 26 NaHCO_3_, 2 CaCl_2_, 1 MgCl_2_ and 10 glucose (pH=7.4, 315mOsm/l) in room temperature. In some cases, technical issues did not allow for electrophysiological recordings, therefore, the number of mice actually used and included in the electrophysiology results were fewer compared to the number of mice used for the behavioral experiments.

The extracellular recording electrode, filled with NaCl (2M), was placed in layer II of the PFC or the stratum radiatum of the CA1 hippocampal region. The platinum/iridium metal microelectrode (Harvard apparatus, Cambridge, UK) was also placed within the upper layers of the PFC or the stratum radiatum layer of the CA1 region of the hippocampus, about 300μm away from the recording electrode, and was used in order to evoke fEPSPs. Responses were amplified using an EXT-02F amplifier (National Instruments), digitized using the ITC-18 board (Instrutech, Inc) on a PC using custom-made procedures in IgorPro (Wavemetrics, Inc, Lake Oswego, OR, USA). Data were acquired and analyzed using custom-written procedures in IgorPro software (Wavemetrics, Inc, Lake Oswego, OR, USA).

The electrical stimulus consisted of a single square waveform of 100 μsec duration given at intensities of 0.1-0.3 mA generated by a stimulator equipped with a stimulus isolation unit (World Precision Instruments, Inc). The fEPSP amplitude was measured from the minimum value of the voltage response compared to the baseline value prior to stimulation. The fEPSP slope was measured from the point that the trace intersected 0mV until 1msec after. Both parameters were monitored in real-time in every experiment. A stimulus-response curve was then determined using stimulation intensities between 0.1-0.3 mA, with a 0.1 mA step. For each different intensity level, two traces were acquired and averaged. Baseline stimulation parameters were selected to evoke a response of 1mV. For the LTP experiments in the PFC, baseline responses were acquired for 10 minutes, then three 1second tetanic stimuli (100Hz) with an inter-stimulus interval of 20 seconds were applied and finally responses were acquired for 50 minutes post-tetanus. For the LTP experiments in the HPC, baseline responses were acquired for 10 minutes, then theta-burst stimulation was applied; 5 pulses at 100Hz every 200ms (theta-rhythm). This stimulation was repeated 3 times with an inter-stimulus interval of 20 seconds. Synaptic responses were normalized to the average 10 minutes pre-stimulus (tetanus or theta-burst) in the PFC and HPC, respectively.

### Statistical Analysis

Data analysis was performed with Microsoft Excel. The data were tested for the presence of outliers but none were identified. The data were first tested for normality using the Kolmogorov-Smirnov test. One-way or repeated measures ANOVA were performed depending on the experiment. Statistical analysis was performed in GraphPad. Data are presented as mean ± standard error of mean (SEM) along with dot plots.

## 3. Results

### Effects of WMT on cognitive flexibility

In the first experiment, 28 adult female mice were used. Mice in the non-adaptive (n=10) group performed the alternating task without any delays, while mice in the fully adaptive (n=10) group underwent the delayed alternation task in the T-maze for 9 days. Mice in the naive group (n=8) remained in their home cage. All the groups subsequently performed the AST task. In the AST task, one-way ANOVA analyses revealed non-significant effects of training in number of trials to reach criterion on the SD, CD, CDR, ID-1, ID-2, ID-2R and ED stages of the task (Figure 1). However, statistically significant differences were found for latency to make a choice between the naïve and adaptive groups in the CD (p=0.03, Tukey’s post-hoc test), the ID-II (p=0.01, Tukey’s post-hoc test) and ID-IIR (p=0.01, Tukey’s post-hoc test) (Figure 2) stages of the AST task. In all three stages of the task post hoc analyses showed that the adaptive group needed less time to choose the bowl in comparison with the naive group.

**Figure 1.**
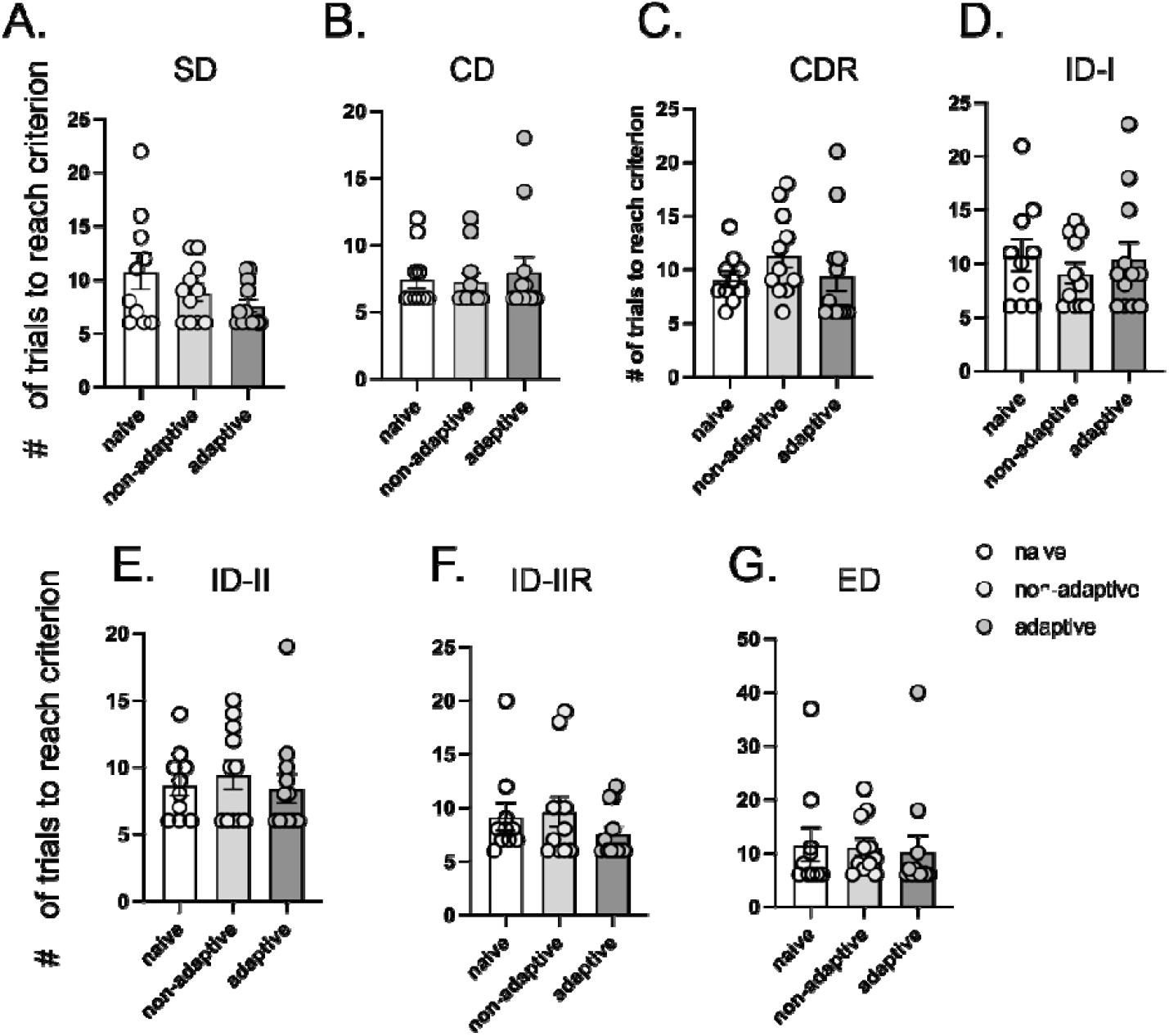
WMT did not affect performance in AST. Graphs show the # of trials required to reach criterion for the different phases of the task and specifically, (A) Simple Discrimination phase (F_(2,30)_=2.26, p=0.12), (B) Complex discrimination phase (F_(2,30)_=0.18, p=0.83), (C) Complex discrimination reversal phase (F_(2,30)_=1.02, p=0.37), (D) Intra-dimensional shift I phase (F_(2,30)_=0.4, p=0.67), (E) Intra-dimensional shift II phase (F_(2,30)_=0.27, p=0.75), (F) Intra-dimensional shift II reversal phase, (F_(2,30)_=0.93, p=0.41), (G) Extra-dimensional shift phase (F_(2,30)_=0.06, p=0.94). F and p values are calculated from one-way ANOVA analysis.

**Figure 2.**
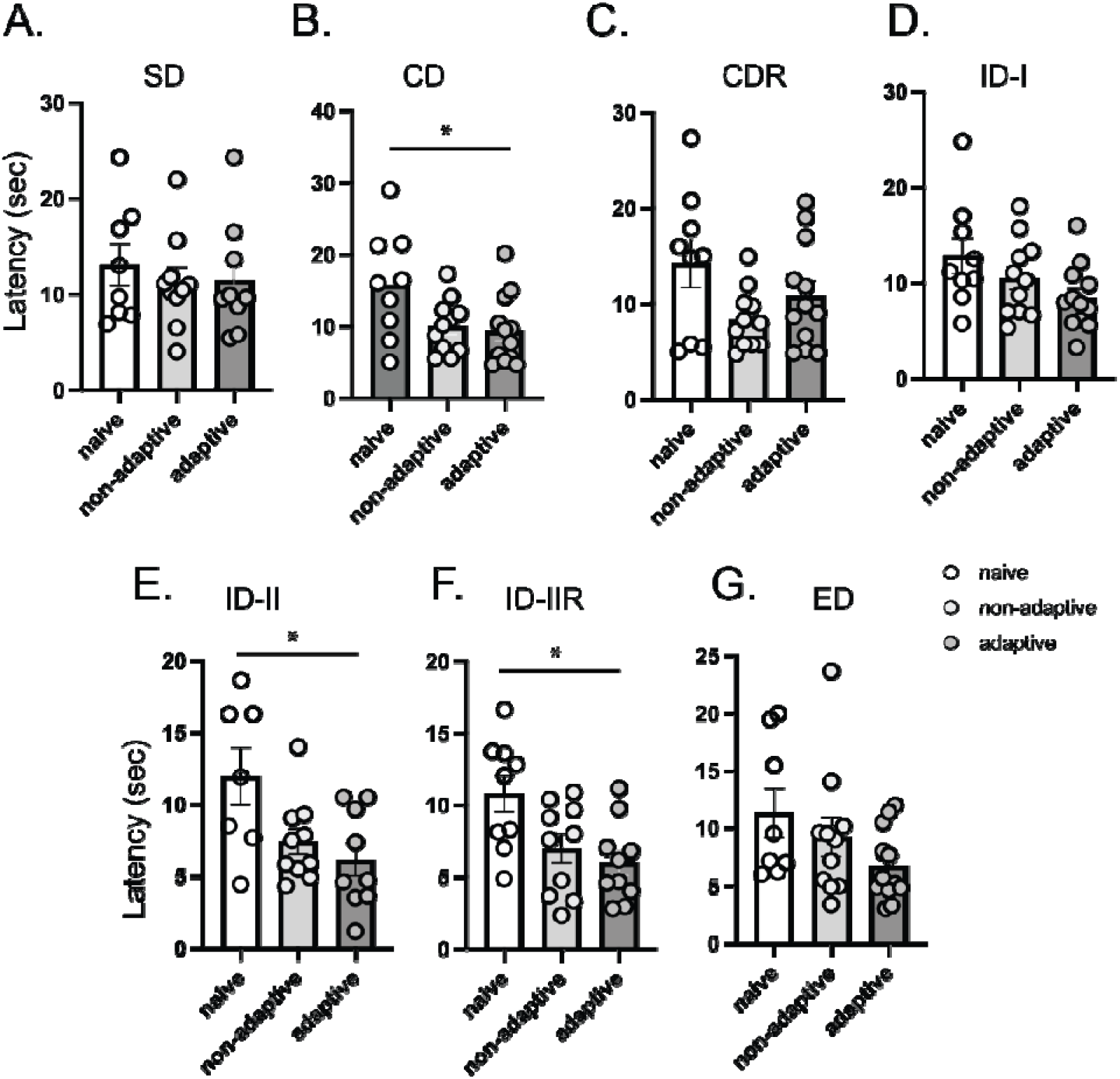
WMT reduced latency in AST. Graphs show the latency to first reach the bowl, and specifically, (A) Simple Discrimination phase (F_(2,30)_=0.26, p=0.77), (B) Complex discrimination phase (F_(2,30)_=4.08, p=0.02), (C) Complex discrimination reversal phase (F_(2,30)_=2.73, p=0.08), (D) Intra-dimensional shift I phase (F_(2,30)_=2.7, p=0.08), (E) Intra-dimensional shift II phase (F_(2,30)_=4.83, p=0.01), (F) Intra-dimensional shift II reversal phase, (F_(2,30)_=5.37, p=0.01), (G) Extra-dimensional shift phase (F_(2,30)_=2.15, p=0.13). F and p values are calculated from one-way ANOVA analysis.

### Effects of WMT on dendritic spine density in the PFC and HPC

Following the AST, Golgi-Cox staining was performed in order to visualise the dendritic spines. First, we compared the three groups for the dendritic spine density of secondary dendrites of PFC pyramidal neurons and identified no statistically significant differences (Figure 3a-c, h). In addition, dendritic spine density was also investigated in the secondary dendrites of the CA1 region of the HPC (Figure 2d-f, g). Here, we identified statistically significant differences in the total and mature dendritic spine density, but not in the stubby dendritic spine density. Specifically, the fully adaptive group presented an increased dendritic spine density in comparison with naive group.

**Figure 3.**
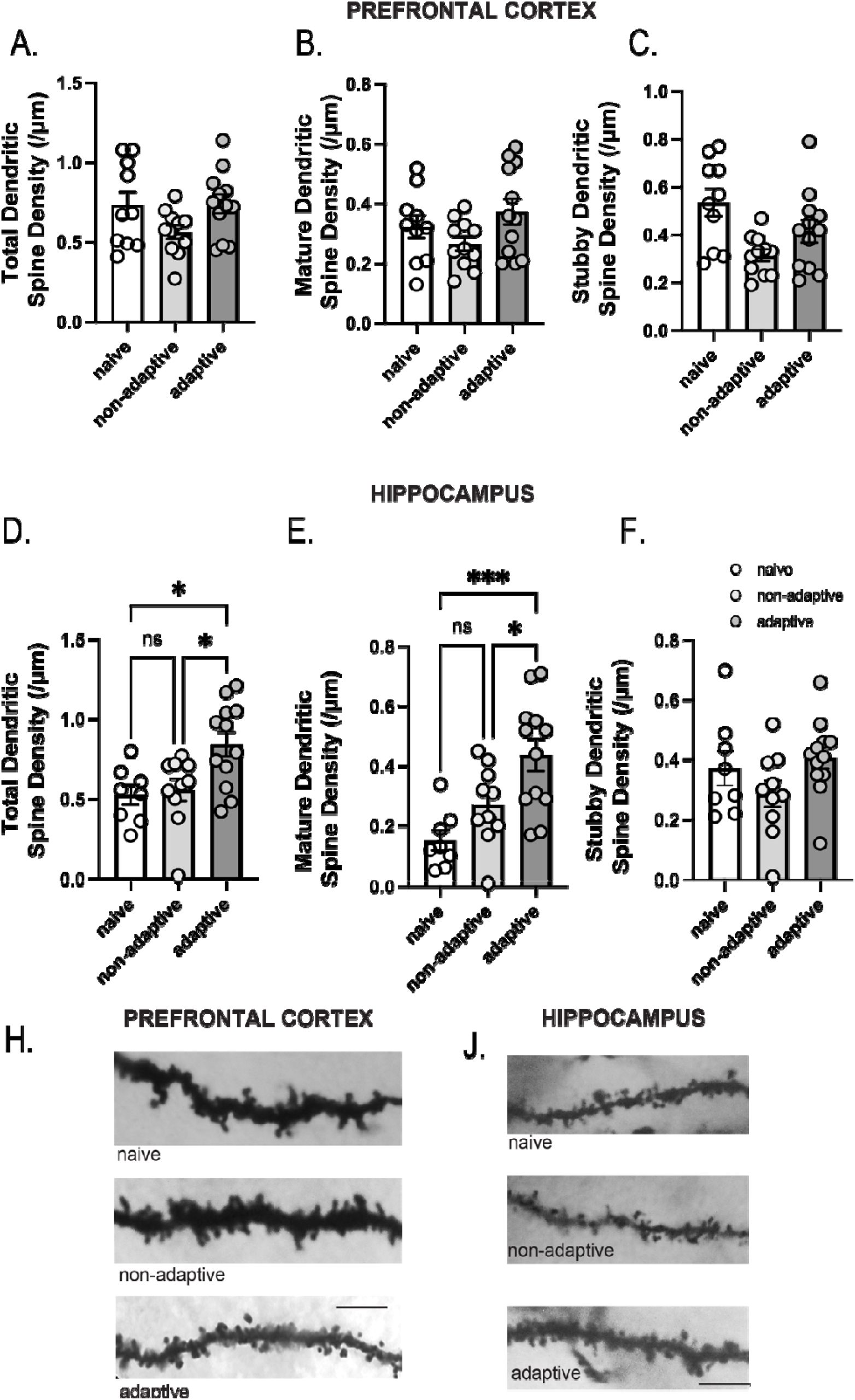
WMT did not have an effect of dendritic spine density in the PFC but increased dendritic spine density in HPC. Graphs showing that there is no difference among the three groups (naïve, non-adaptive and adaptive) in the total dendritic spine density (F_(2,30)_=2.52, p=0.09) (A) and the mature dendritic spine density (B) (F_(2,30)_=2.22, p=0.12) in the prefrontal cortex. There is a significant difference in stubby dendritic spine density (F_(2,30)_=5.68, p=0.008) among the three groups, and specifically a reduction in the non-adaptive group compared to the naïve group (Tukey’s test, p=0.0057). For the hippocampus, there is a significant difference among the three groups in the total dendritic spine density (D) (F_(2,27)_=6.15, p=0.0063) and specifically, the total dendritic spine density is increased in the adaptive compared to the naive and non-adaptive groups (Tukey’s test, p=0.0147 and p=0.0192, respectively). (E) There is a significant difference among the three groups in the mature dendritic spine density (F_(2,27)_=9.178, p=0.0009) and specifically, the mature dendritic spine density is increased in the adaptive compared to the naive and non-adaptive groups (Tukey’s test, p=0.0008 and p=0.0379, respectively). (F) There is no significant difference among the three groups in the stubby spine density in the hippocampus (F_(2,30)_=2.018, p=0.15).

### Effects of WMT on synaptic response and synaptic plasticity in the PFC and HPC

In a second cohort, mice were split into 3 training groups as above. Ex vivo electrophysiological recordings were performed to evaluate the synaptic response and synaptic plasticity in the PFC and HPC. With regards to the PFC, the adaptive group included n=5 mice, the non-adaptive n=5 and the naive n=6. In the HPC, the adaptive group comprised n=5 mice, the non-adaptive n=6 and the naive n=6 mice. No significant differences were recorded in the evoked fEPSPs in response to increasing stimulation in both the PFC and HPC (Figure 4). Furthermore, we examined the induction of LTP in the PFC and HPC in mice that were previously subjected to WMT. In the PFC, tetanic stimulation resulted in a significant increase in the synaptic response in all three groups, without any significant difference among them (Figure 5A). In the HPC, theta-burst stimulation resulted in synaptic potentiation in all three groups. However, the enhancement of synaptic potentiation in the adaptive groups was significantly higher compared to the naïve and non-adaptive groups (Figure 5B), supporting the involvement of HPC in WMT.

**Figure 4.**
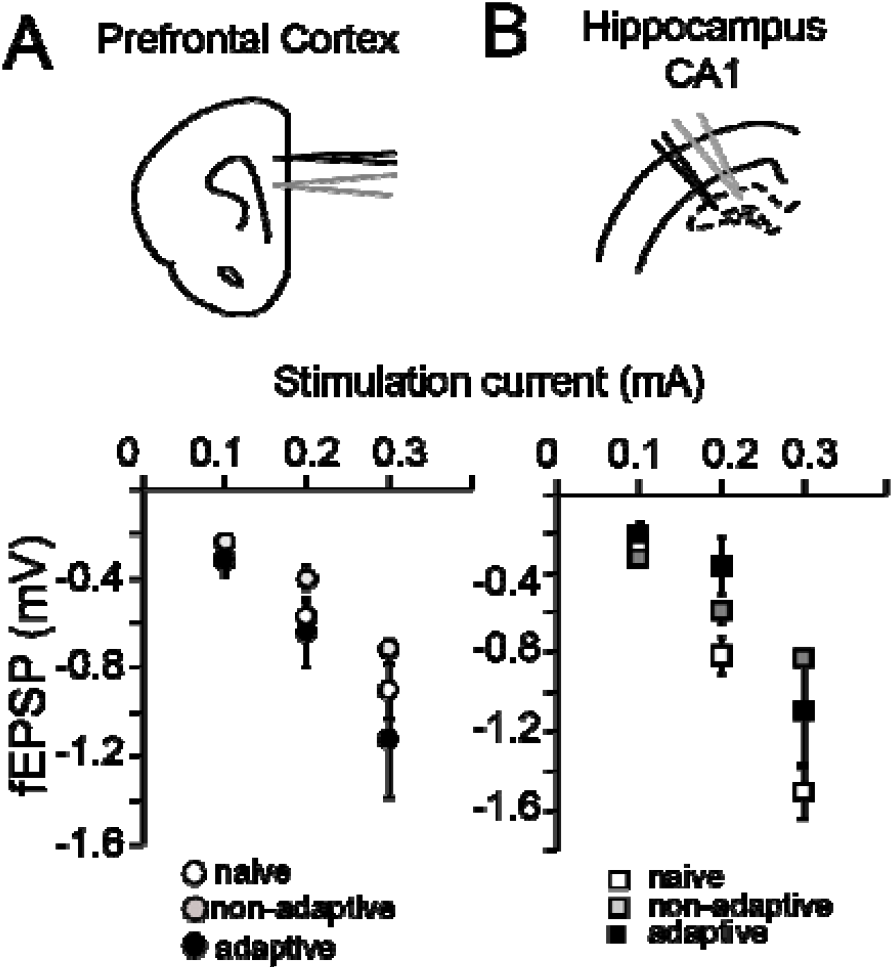
WMT does not alter the evoked fEPSPs in the PFC and HPC of female mice. Synaptic transmission in the PFC and the HPC, in the naïve, non-adaptive, and adaptive groups. (A) (Top) Electrode layout in layer II of the PFC. (Bottom) Graph showing that no difference in the fEPSP amplitude in response to increasing current stimulation in the PFC between the three training groups, in the range of 0,1 mA-0,3 mA. (B) (Top) Electrode layout in the CA1 subfield of the HPC. (Bottom) Graph showing that there is no significant difference in the fEPSP amplitude in the response to increasing current stimulation in the HPC between the three different groups, in the range of 0,1 mA-0,3 mA.

**Figure 5.**
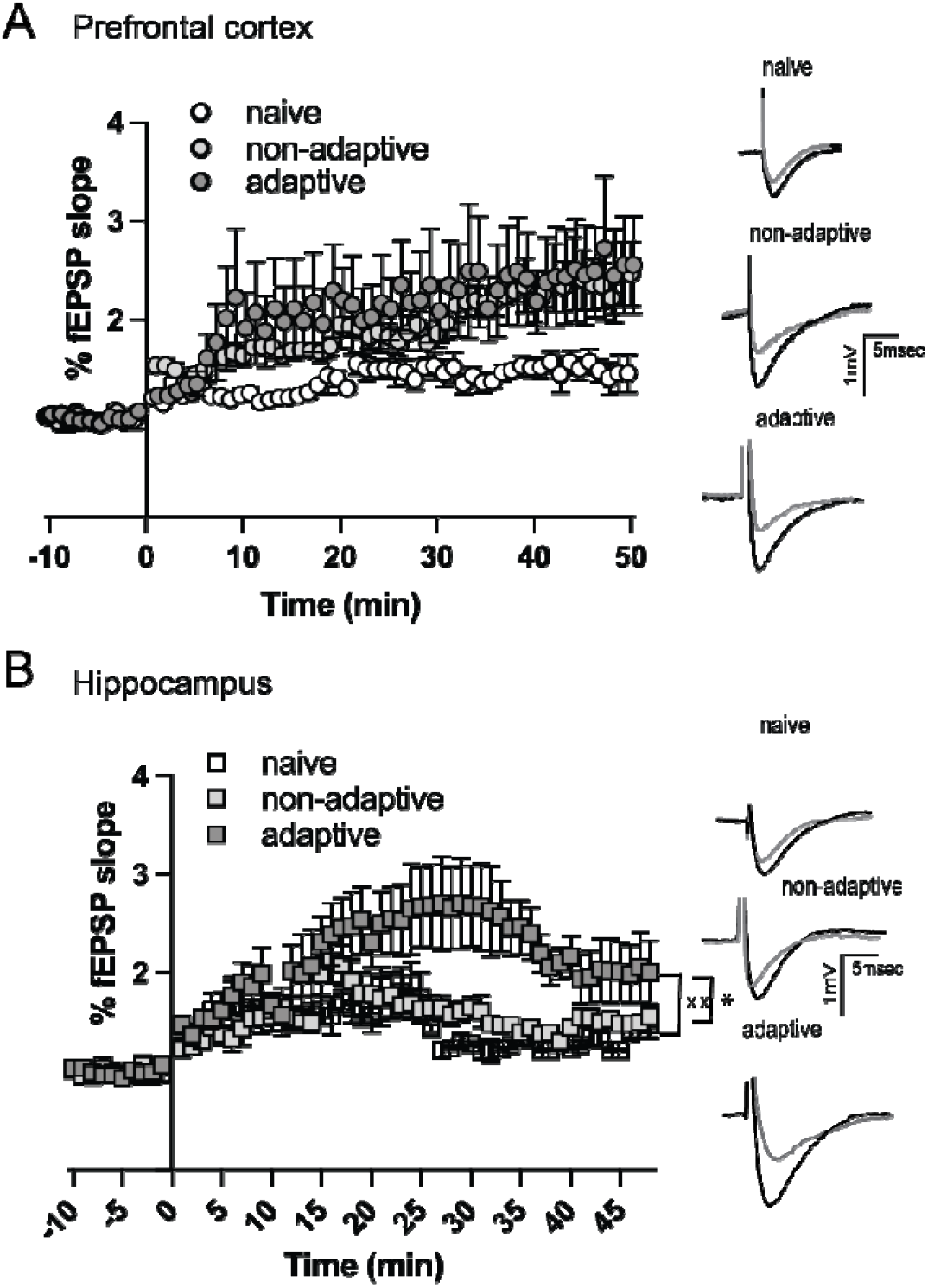
Long-term potentiation in the PFC and the HPC of naïve, non-adaptive, and adaptive groups. (A) Graph (left) and representative traces (right) showing the potentiation of the fEPSP following tetanic stimulation in the PFC of naïve, non-adaptive and adaptive adult female mice. Repeated measures ANOVA did not identify any significant difference among the three training groups (F_(2,14)_=3.1, p=0.15) (B) Graph (left) and representative traces (right) showing the potentiation of the fEPSP following theta-burst stimulation in the HPC of naïve, non-adaptive and adaptive adult female mice. Repeated measures ANOVA identified a significant difference among the three training groups (F_(2,14)_=8.5, p=0.0037). In particular, there was a significant difference between the naïve vs the adaptive group (p=0.0068) and between the non-adaptive and the adaptive groups (p=0.013) (Sidak’s multiple comparison test).

## Discussion

In this study, we investigated the effects of WMT on cognitive flexibility, dendritic spine density and synaptic plasticity in the PFC and HPC of female adult mice. We found that WMT did not affect the performance in the AST task but reduced the decision latency in some stages of the task. Furthermore, WMT did not affect dendritic spine density or LTP in the PFC, however, WMT increased dendritic spine density and LTP in the HPC of female mice.

### Comparing the effects of WMT in male and female mice

In our previous study, we found that WMT in adult male mice improved performance in the extradimentional shift of the AST task, increased density in pyramidal neurons in the PFC and HPC and increased LTP in the PFC (Stavroulaki et al., 2021). On the contrary, we have not identified a beneficial effect of WMT on cognitive flexibility in female mice, although there was a significant reduction on decision latency in the CD, ID-II and ID-IIR stages of the task.

The reduced decision latency could indicate that WMT in females also enhances the function of neuronal circuits involved in information processing (Robbins, 2002; Scheggia et al., 2014). Thus, while not translated in performance improvement, WMT does seem to have some beneficial effect on cognitive flexiblity in females, as well.

Furthermore, in our previous study using male mice, we identified enhanced dendritic spine density both in the PFC and HPC as well as increased LTP only in the PFC (Stavroulaki et al., 2021). In the currently study using female mice, we identified increased dendritic spine density and LTP in the HPC, but no changes in the PFC. These results suggest that in males, PFC function can be enhaced with WMT while in females HPC function is improved. These differences between males and females indicate sex differences in the enhancement of synaptic plasticity, in agreement with previous reports (Koss and Frick, 2017; Safari et al., 2021).

### Sex differences observed in WM and cognitive flexibility

We have not identified significant baseline differences between male and female mice in the delayed alternation task (Chalkiadaki et al., 2019), the attention set-shifting task (this work and Stavroulaki et al., 2021), as well as other behavioral tasks that depend either on the PFC or HPC, such as novel object recognition, object to place and temporal order object recognition (Plataki et al., 2021). A recent, independent study also did not identify any changes in the memory-related component of object recognition tasks and in working memory, using the water maze (Melgar-Locatelli et al., 2024). These results suggest that there are no baseline differences between male and female C57Bl/6 mice that could account for sex differences observed in the effects of WMT. Other studies using primarily rats have identified improved performance in males compared to females in spatial memory and WM using the radial arm maze, as well as in object location memory tested 24 hours after first object exploration ((Koss and Frick, 2017). However, superior performance in spatial WM has also been observed in female rats (Bimonte et al., 2000). Therefore, it seems that sex differences in cognitive function depends on the species, strain as well as task parameters (Koss and Frick, 2017). Therefore, it is possible that sex differences in the effects of neuromodulators could account for the differential effects of WMT on cognitive flexibility, PFC and HPC function (Li et al., 2016; Velli et al., 2021).

### Sex differences in dendritic spine density and LTP

A few studies have investigated baseline sex differences in dendritic spine density and LTP in the HPC. Most studies do not identify significant differences in dendritic spine density in the CA1 region, although modulation of estrogen levels can affect dendritic spine densities (Koss and Frick, 2017). Specifically, it seems that in the female HPC, LTP has an increased threshold

Regarding the PFC, dendritic spine density in the females (this study) is not different compared to the male study (Stavroulaki et al., 2021).

LTP in the CA1 hippocampal region has exhibited sex differences that could be attributed to the estrous cycle of the females (Koss and Frick, 2017). Specifically, it seems that LTP in females has a higher threshold and activation of membrane estrogen receptors E2 can enhance LTP(Gall et al., 2023). However, no evidence that LTP is different between male and female mice or that it is affected by the estrous cycle has been found for the prefrontal cortex (Velli et al., 2022).

### Female PFC function seems to be more resistant to change compared to male PFC

No studies have examined the possibility to enhance cognitive function in female mice, to the best of our knowledge, however, there are few studies that have examined cognitive function in response to different manipulations that usually result in impairments. Acute stress reduced performance in the temporal order object recognition task (TOR) in adult male, but not female mice (Velli et al., 2021). WM is significantly impaired in male, but not female mice, of the schizophrenia MAM model of schizophrenia (Chalkiadaki et al., 2019). Male mice with hypofunction in the dysbindin gene, a schizophrenia risk factor gene, exhibited impaired performance in the TOR task, but not female mice (Geraci et al., 2024). It is possible that differential modulation of neuromodulator systems could contribute to these difference between males and females (Sannino et al., 2015; Li et al., 2016). However, that may not always be the case, as chronic stress has similar deteriorating effects in both males and females (Anderson et al., 2019). Therefore, it seems that PFC function in females is more resistant to change (improvement or impairment) compared to males.

## Acknowledgements

This work was supported by the Special Accounts for Research of the University of Crete and the Marie Curie RISE neuronsXnets grants.

